# Associated factors and spatial patterns of the epidemic sporotrichosis in a high density human populated area: A cross-sectional study from 2016 to 2018

**DOI:** 10.1101/693085

**Authors:** Lívian Otávio Lecca, Marcelo Teixeira Paiva, Camila Stefanie Fonseca de Oliveira, Maria Helena Franco Morais, Maria Isabel de Azevedo, Camila de Valgas e Bastos, Kelly Moura Keller, Roselene Ecco, Márcia Regina Silva Alves, Graziella Coelho Tavares Pais, Lauranne Alves Salvato, Gustavo de Morais Donancio Xaulim, David Soeiro Barbosa, Silvana Tecles Brandão, Danielle Ferreira de Magalhães Soares

## Abstract

We carried out an epidemiological characterization of human and feline sporotrichosis, between 2016 and 2018, in a high density-populated area in Brazil. Professionals were trained to identify suspected cats and notified vets to interview the owners and collect swabs of the wounds from these animals. Mycological cultures were performed, and colonies identified for Spotrothrix spp. Subsequently, data regarding the outcome from suspected animals were collected. Confirmed cases of human sporotrichosis (56) were also counted and analysed for spatial distribution. Regions with highest prevalence of feline sporotrichosis, had greater frequencies of both human and feline cases. 118 (77.63%) animals were positive. Animals that lived only partially at home were 3.02 times more likely of being positive (OR 3.02, CI 95% 1,96-10,43). The prevalence of feline sporotrichosis was 8.36 ‰ (CI 95%, 5.38 - 9.55 ‰). There was no statistically significant association between environmental variables and positive diagnosis, corroborating the hypothesis that direct transmission by infected cats plays a greater role in the occurrence and continuous outbreaks of sporotrichosis in Brazil. Among the positive animals, 61.90% (CI 95% 58.95 - 64.96) died, being 6.30 times more likely to die than negative animals (p< 0.05, OR 6.30, CI 95% 2,79-14,42). The lethality rate was 55.08% in cats (CI 95% 49.20 - 51.15). The mortality for sporotrichosis was 4.6 ‰ cats (CI 95% 3.4 - 6 ‰). Only 7.62% (CI 95% 7.12 - 8.16) positive cats were treated and cured. Among dead positive animals, 29.23% were inappropriately discarded. This is the first report on the epidemic of sporotrichosis in Minas Gerais, Brazil. The free offer for treatment and veterinary care to these animals should be taken into consideration, as well as the collection and incineration of the dead ones, as measures of public health, followed by the guidance and care for the human patient.

## Introduction

Sporotrichosis is a cosmopolitan zoonotic disease caused by fungi of the complex *Sporothrix schenckii*, with high relevance in Brazil. Since the first reports of its transmission to humans through contact with infected cats in the 1990s, the number of severe cases of the disease in humans has been increasing. Therefore, this zoonosis has become relevant to public health [1–4]. The infected cats carry the fungus in their injured sites, oral cavities, and nails, and the transmission is made by scratches, bites, and contact with the wound exudates [5–7].

Cases of the disease in cats and humans have been described in the northern, eastern, and southern states of the country [7–10]. In the state of Minas Gerais, there are no descriptions in literature about any epidemiological situation, although it shares boundaries with the state of Rio de Janeiro, a hyperendemic region for sporotrichosis. Only one study has reported the presence of *Sporothrix brasiliensis* in a human case and ten domestic cats in the state [7].

Considering infected cats as potential transmitters of the disease to humans and the increasing territorial expansion of sporotrichosis in the country, it is necessary to characterize the epidemiological situation of the disease in areas with potential spread of the infection.

Accordingly, the present study aimed to produce the epidemiological characterization of human and feline sporotrichosis in a high density-populated area located in the municipality of Belo Horizonte (Minas Gerais, Brazil), based on the first reports of sporotrichosis. This included the analysis of the spatial distribution, associated factors, and the identification of the most common outcomes of such cases in cats, in the period from August 2016 to June 2018.

## Material and Methods

### Ethics approval

This study was approved by the Universidade Federal de Minas Gerais, Comitê de Ética em Pesquisa da Universidade Federal de Minas Gerais (Committee for Ethics in Research of the Federal University of Minas Gerais) (number CAAE – 67149517.5.0000.5149).

### Type and area of study

We conducted a cross-sectional epidemiological study based on observation, laboratory diagnosis of sporotrichosis in domestic cats, and analysis of factors related to the disease and the respective outcomes. In addition, we incorporated the spatial analysis of cases in cats and humans, in the Barreiro region, Belo Horizonte, Minas Gerais, between 2016 and 2018.

Belo Horizonte, the capital of the state of Minas Gerais, is located in southeastern Brazil, and it is the sixth most populous city of the country. It has a geographic extension of 331.401 km^2^ and a population density of 7,167 inhabitants per km^2^. The city is divided into nine regional administrative units. Of these regions, Barreiro (an area of study), which is subdivided into 20 areas, is covered with Health Centers. The selection of this study area was determined based on the fact that it has the largest population of cats compared to other regions of the municipality, and because the first reports of suspected sporotrichosis in the municipality were reported from Barreiro.

### Selection, samples collection, and data

Between August 2016 and June 2018, the Escola de Veterinária da Universidade Federal de Minas Gerais (School of Veterinary Medicine of the Universidade Federal de Minas Gerais) (EV/UFMG), in partnership with Diretoria de Zoonoses (Council on Zoonosis) (DZ/SMSA), in the Municipality of Belo Horizonte, received samples and cadavers of cats suspected of sporotrichosis, from the county of Barreiro.

The public health professionals in that region were then trained to identify nodular and ulcerative skin lesions in cats, potentially caused by sporotrichosis. After the suspected animals were identified, the vets of the DZ/SMSA were notified to interview the owners and collect samples from the animals. The period of search for suspected cases was extended until the end of the study (2016 and 2018), and the visits of veterinarians to the notified homes continued for the whole period.

The owners of the suspect animal answered a semi-structured questionnaire about the characteristics of the symptomatic animals and the home itself, to identify possible factors associated with the occurrence of sporotrichosis and conduct of those responsible for the animals face to that suspicion.

Swabs of the wounds of the suspected cats were obtained during the home visits and the corpses of the dead cats were forwarded by the DZ/SMSA to the Laboratory of Veterinary Pathology of the EV/UFMG.

Mycological cultures were then performed in the Laboratory of Mycology and Mycotoxins (LAMICO), of the School of Veterinary Medicine of the UFMG, following the cultivation into a solid media on Petri plates. Each sample was incubated in the media BHI/36 °C (BD, USA) and SDA/25 °C (Kasvi, Brazil). Colonies were identified by their macro and micromorphology [11,12].

From April to June 2018, the veterinarians of DZ/SMSA monitored the study area to identify cats suspected of sporotrichosis. The veterinarians investigated and collected data including as suspicious cats remained at their homes, progression of the clinical signs, completion of treatment after confirmation of the disease and destination of the corpses in the event of death.

In addition, confirmed cases (with positive culture) of human sporotrichosis occurred in this same period were also counted and analysed for spatial distribution.

### Preparation and analysis of data

The data from LAMICO and DZ/SMSA were entered in spreadsheets, and were grouped according to the four variables - Demographic, Environmental, Diagnosis, and Outcome. In cases where inconsistency was found in a record without the possibility of verification and correction, the record was excluded from the statistical analysis. For the analyses of environmental factors, lack of data on the environmental variables was considered one exclusion factor.

The descriptive analysis of the data included the calculation of averages, frequency of distributions, coefficient of prevalence, frequency of positivity, coefficient of mortality, and lethality rate, with their respective confidence intervals at 95% (CI 95%), considering the local feline census in 2018 as the total population.

The association analyses of the qualitative variables with the diagnosis was carried out through the chi-squared test (CI 95%; p < 0.05) using the software Stata 14.0. Furthermore, Spearman correlation analysis (CI 95%; p < 0.05) was performed between the variables of population density and prevalence of the disease in felines, using the packages *stats* and *ggpubr* of the software R.

For the analysis of the spatial distribution, the cases were geo-referenced based on the addresses of the households visited. Maps of population density, prevalence of sporotrichosis cases, and Kernel density were obtained using the QGIS^®^ software 2.18.

## Results and Discussion

### Zoonotic sporotrichosis

Between 2016 and 2018, 56 human cases of sporotrichosis (Fig 1) were recorded in the study area. The regions where the rate of prevalence of feline sporotrichosis was above 6.7%, were included 69.6% (39/56) of human cases and 66.4% (101/152) of feline cases (Fig 2). The areas of high prevalence of feline sporotrichosis, as highlighted in Fig 1, the calculation jointly accounted 42.8% of human cases. In two regions where no positive feline cases were found, human cases were recorded. This may indicate the presence of non-identified feline cases in those areas or that no animal transmission to humans occurred in the residence.

**Fig 1 -.**
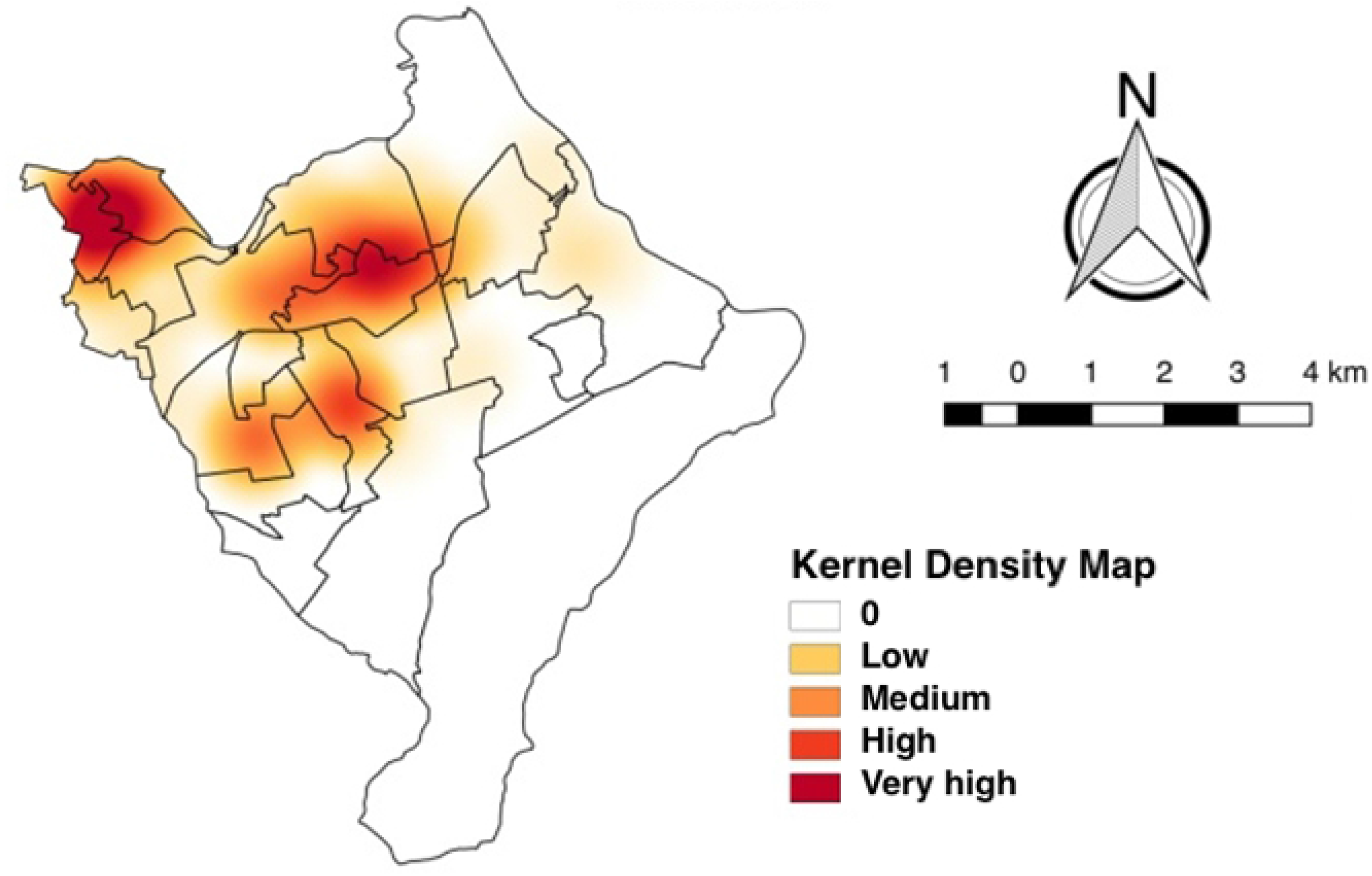
Spatial distribution of sporotrichosis cases in humans and cats in the county of Barreiro, Belo Horizonte, 2016 to 2018.

**Fig 2 -.**
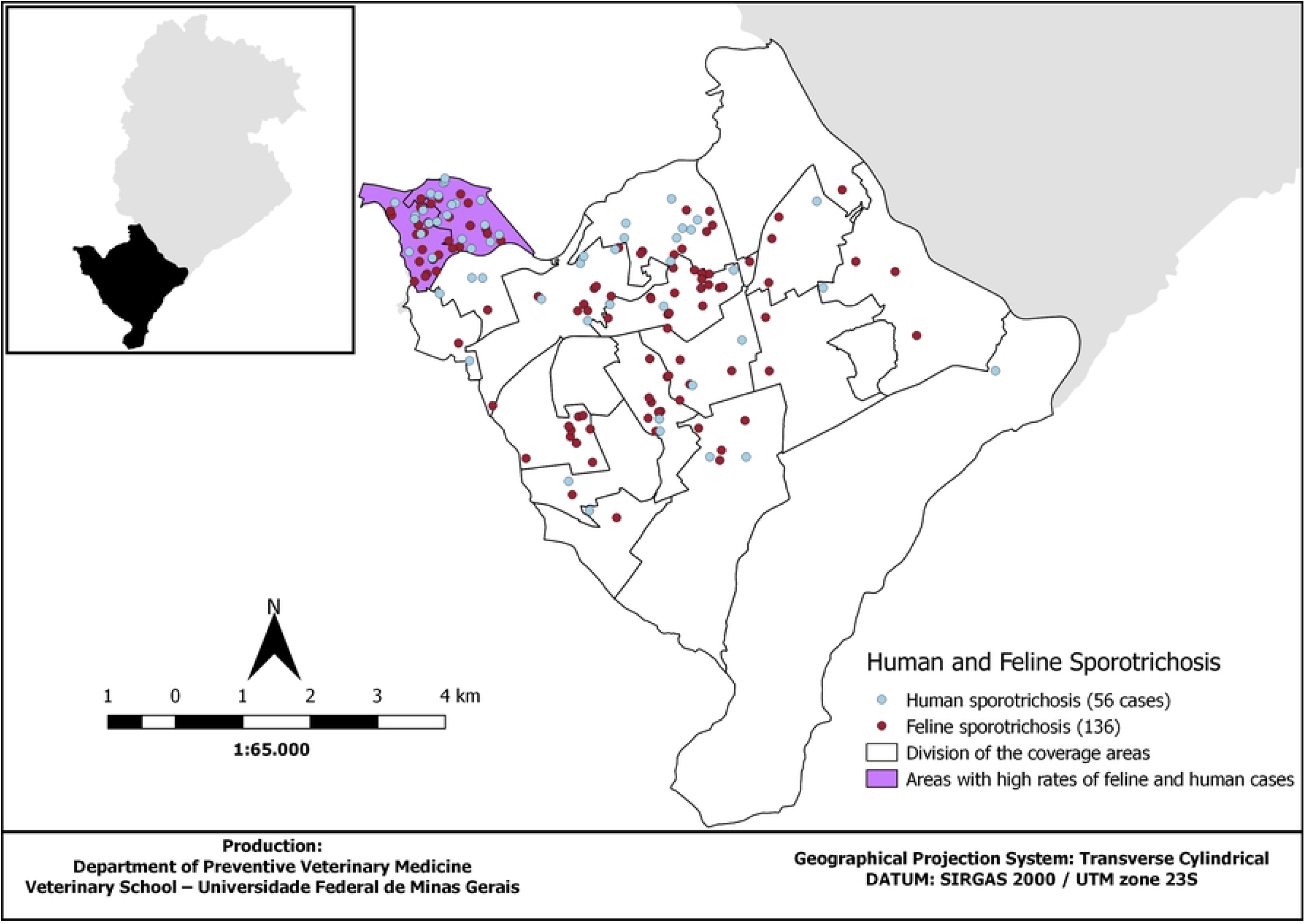
Spatial distribution of human cases of sporotrichosis in relation to the prevalence of feline sporotrichosis per area of coverage in the county of Barreiro, Belo Horizonte, 2016 to 2018.

### Feline sporotrichosis

During the study period, samples were collected from 152 cats for laboratory diagnosis. One hundred and thirty seven (90.13%) cats were identified as suspect and the corpses of 15 (9.87%) dead cats were sent for necropsy. Sixteen animals with samples collected *in vivo* were excluded from statistical analyses of association, due to lack of data. Information regarding the 15 corpses was not included in the analysis of environmental variables, because questionnaires were not answered by the cat owners.

The sample was divided into two groups: positive and negative. The inclusion in each category was based on epidemiological, clinical, and/or laboratory criteria (Fig 3). Among the samples with positive isolation, one was sent for molecular analysis, which was confirmed as *S. brasiliensis* through molecular diagnosis [13]. For the samples of corpses, diagnosis was based only in laboratorial results. Thus, 118 (77.63%) animals were considered positive while 34 (22.37%) animals were considered negative (Table 1).

**Fig 3.**
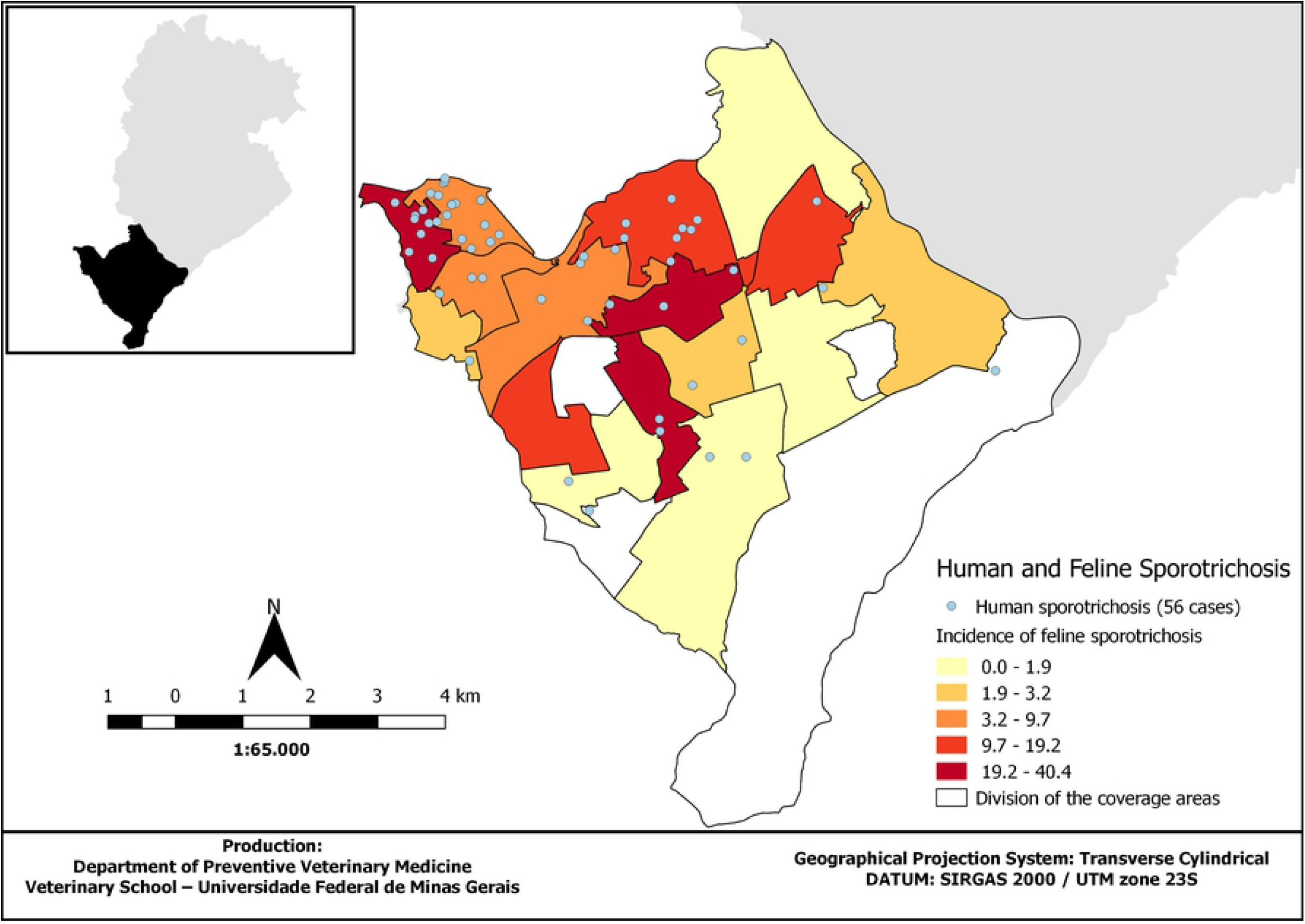
Criteria for the diagnosis of feline sporotrichosis in the county of Barreiro, Belo Horizonte, Minas Gerais, August 2016 to June 2018.

**Table 1 -.** Distribution of household cats in relation to the diagnosis of sporotrichosis in the county of Barreiro, Belo Horizonte, Minas Gerais, August 2016 to June 2018.

|  |  |
| --- | --- |
| **Field collections of suspect cats** |  |
| Laboratory positive | 75 (54.74%) |
| Clinical-epidemiological positive with laboratory negative | 16 (11.68%) |
| Clinical-epidemiological positive, going through prior treatment with laboratory negative | 12 (8.76%) |
| Clinical-epidemiological and laboratory negative | 34 (24.82%) |
| Total | 137 (100%) |
| **Corpses of suspected cats** |  |
| Laboratory positive corpses | 15 (100%) |
| Laboratory negative corpses | - |
| Total | 15 (100%) |
| **Final diagnosis** |  |
| Positive | 118 (77.63%) |
| Negative | 34 (22.37%) |
| Total | 152 (100%) |

### Characteristics of the feline population

The distribution of characteristics in the population composed of domestic cats studied was similar between the positives and negatives groups. Among the animals of known age, majority were young adults (aged between 1 and 3 years) (38.97%). There was predominance of animals without a defined breed (WDB) (98.53%), male animals (65.44%), not surgically spayed animals (69.12%), and animals with a lifestyle spent partially at home (68.38%) (S1 Table).

For animals with a positive diagnosis, there was predominance of young animals of known age (1 to 3 years) (33.33%) and males (64.76%). Similar results were obtained in studies on endemic and outbreak regions in Rio de Janeiro, Rio Grande do Sul, and Recife [9,10,14,15]. Majority of the animals was not spayed (68.57%); however, it was not possible to verify a significant association (p > 0.05) between castration and positive diagnosis. Furthermore, in both groups, the proportion of animals that would regularly wander the streets was high (83.3% of non-spayed and 74.08% of spayed animals). Such information shows that, even for those guardians who decided spaying their cats, confinement is not considered an important measure for animal care.

Only the lifestyle presented a statistically significant association with positive diagnosis of sporotrichosis (p < 0.05). Of the animals assessed, 86.76% (118/136) lived only partially at home. These animals were 3.02 times more likely of being diagnosed as positive for sporotrichosis compared to animals strictly confined to their homes (OR 3.02, CI 95% 1,96-10,43). The finding is similar to that obtained in previous studies conducted in southern Brazil [16], which shows a considerable percentage of animals with free access to the street and a low degree of responsible ownership by their guardians. Such behavior exposes the animals to fights, copulation, and contact with sick animals. Additionally, the sick cat may act as a disseminator of the fungus to other animals and to the environment [14,16].

All the variables (young animals, male, non-spayed, and with access to the street) are known risk factors of the disease, according to previously published studies [5,14]. For the improvement of a responsible ownership, it is of great importance to spread the information about this serious disease presenting a risk of infection for the animals and for the families they live with.

As majority of the population included in the study comprised young, non-spayed cats with free access to the street, this homogeneity did not allow the quantification of the effect of these variables on the occurrence of the disease in this population.

### Spatial distribution of feline cases

In relation to the spatial distribution of the sample, 16 (80%, CI 95%, 56.34% - 94.27) of the 20 areas showed at least one positive cat for sporotrichosis. The prevalence found in the county of Barreiro was 8.36 positive cats per one thousand cats (CI 95%, 5.38 - 9.55 per one thousand cats – S2 Table).

The three areas with the highest prevalence were not in the group of areas with higher cat density per household (Fig 4). The cat population density and the prevalence were analyzed with the Shapiro-Wilk test, and the results showed that only population density exhibited normal distribution (p > 0.05). In addition, Spearman correlation test was carried out between such variables, but no statistically significant correlation between them was found (p = 0.09). This result may have occurred because the calculation of density did not include an estimate of the free roaming population, of which there were no available data, and they may contribute to the greatest infection rate in cats with access to the street. Another possibility is that the risk of infection is more related to the cultural habits of the owners, giving cats a free access to the street, than the local density. This is because studies have shown that cats with access to the street may wander across large areas, up to 1.92 ha [17,18].

**Fig 4 -.**
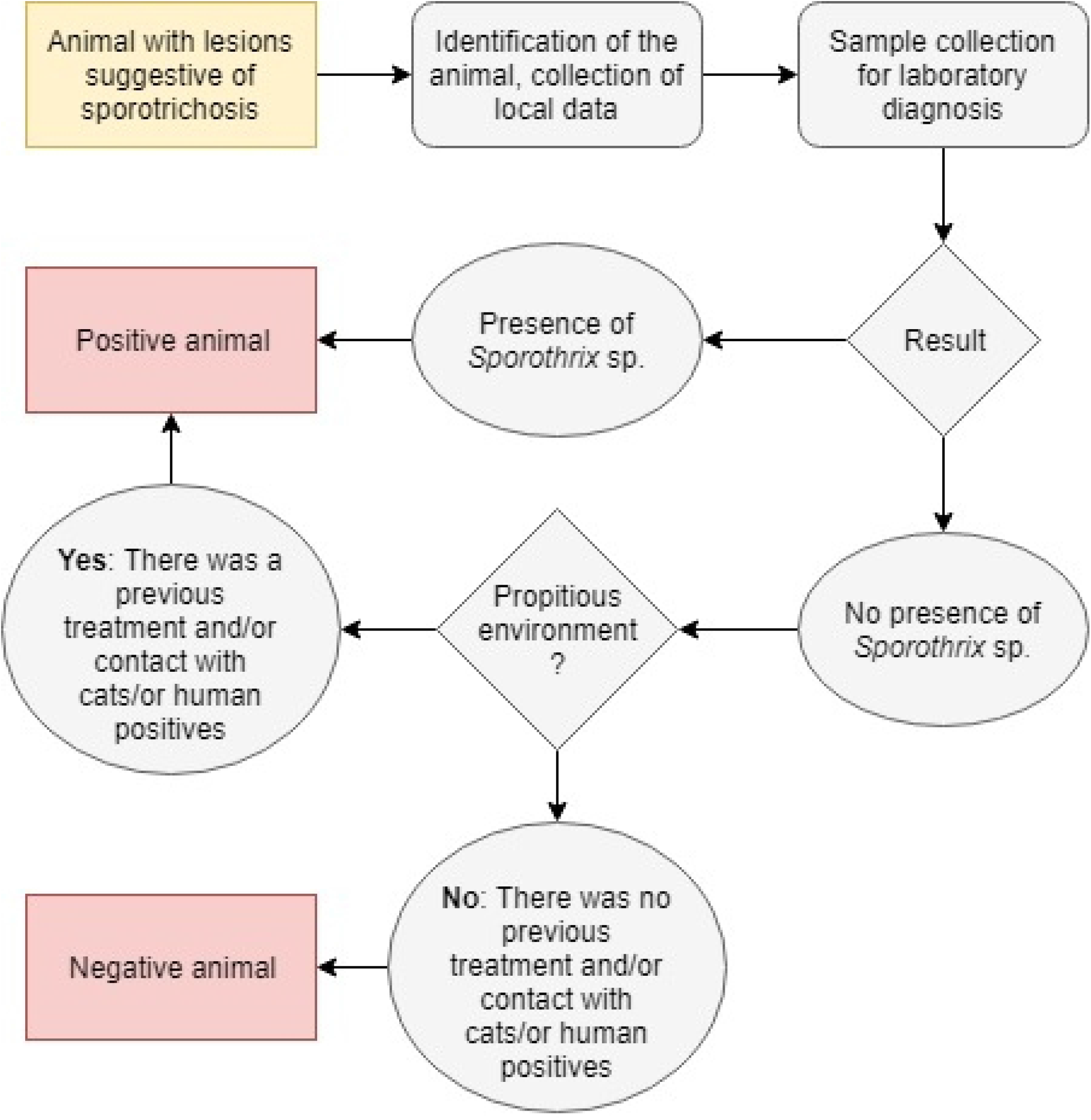
Map of feline population density (cats per 100 residences, A), and the prevalence of sporotrichosis (cases per thousand cats, B) in the county of Barreiro, in Belo Horizonte, 2016 to 2018.

Five areas of coverage amounted to a participation of 71.19% (CI 95% 70.29 - 72.07) of the total number of positive cats. The Kernel analysis showed the existence of three areas where cases were centered (Fig 5). These results may be indicators of areas where actions for the prevention and control of the disease should be prioritized and intensified.

**Fig 5 -.**
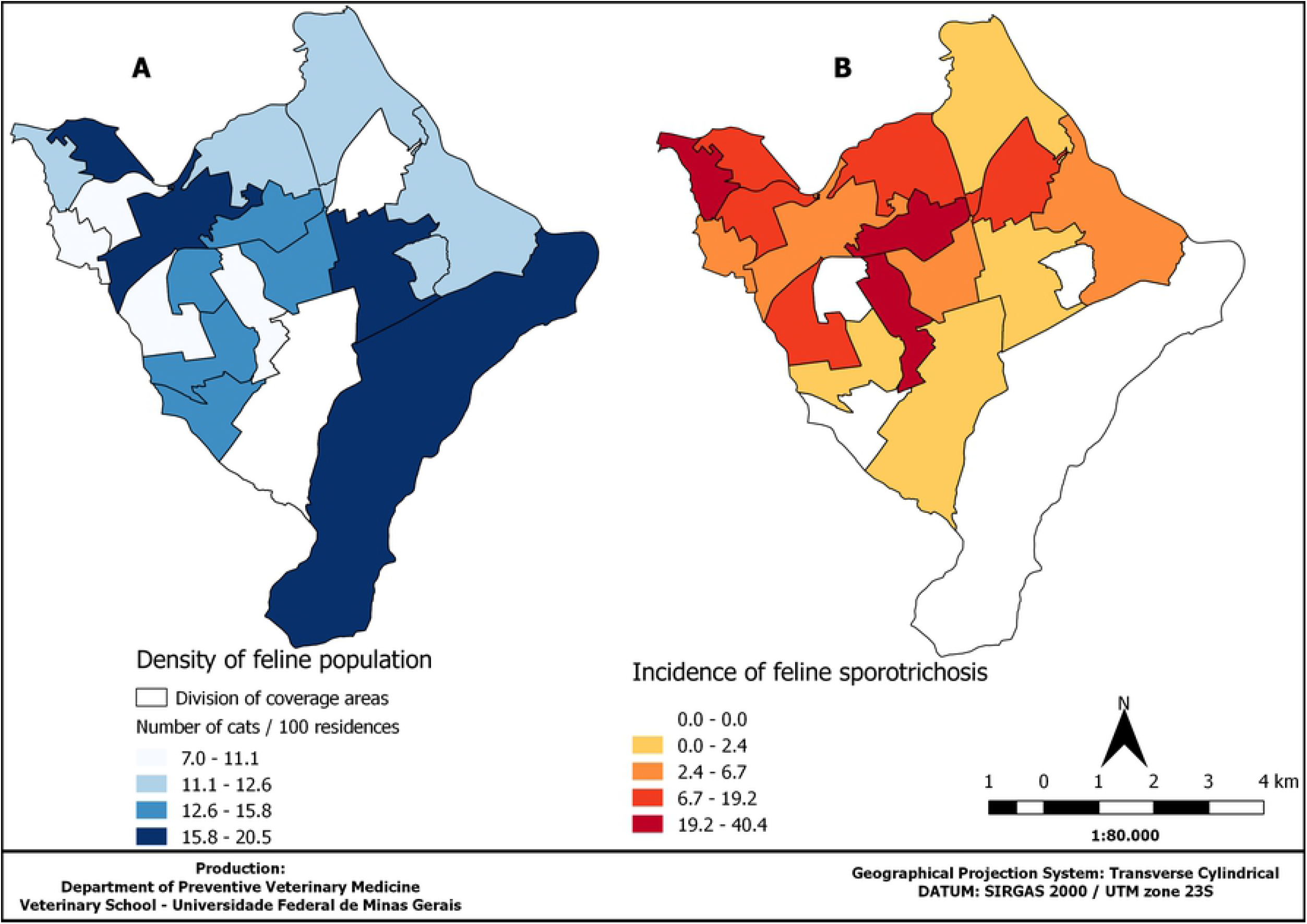
Map of the intensity of sporotrichosis cases in the county of Barreiro, Belo Horizonte, 2016 to 2018.

### Environmental Conditions

The distribution of characteristics related to the homes of the analyzed cats (S3 Table) was similar between positive and negative animals groups, with predominant houses (94.21%, CI 95% 89.52 - 98.16) having a backyard (90.98%, CI 95% 87.18 - 97.05). Organic matter was observed in 49.59% (CI 95% 40.74 - 59.26) of the households. The presence of debris (27.27%, CI 95% 20.28 - 37.27), trees (24.79%, CI 95% 18.02 - 34.54), vegetable gardens (18.18%, CI 95% 12.28 - 27.29), and grass (15.70%, CI 95% 10.07 - 24.19) was less common.

There was no statistically significant association between environmental variables and positive diagnosis for sporotrichosis. It was observed that the characteristics of households in the sample studied, as well as the characteristics of cats, were quite homogeneous, which did not allow the quantification of the association between them and the diagnosis of sporotrichosis.

It is important to emphasize the existence of the only article in literature, which reported that it was possible to isolate *S. brasiliensis* from environmental samples, a more frequent species in the outbreaks taking place in Brazil [19]. These results seem to corroborate the hypothesis that direct transmission by infected cats plays a greater role in the occurrence and continuous outbreaks of sporotrichosis in Brazil [20], although the possibility of a classical infection route in these places should not be discarded.

### Outcome of positive domestic cat cases

It was possible to get data on the outcome of 96.32% (95% CI 91.63 - 98.80, 131/136) of the cats assessed (S4 Table); however, 62.49% (95% CI 53.79 - 70.65, 85/136) were not present at the residence after the period between the first and second visits to the site. Among the positive animals, 61.90% (CI 95% 58.95 - 64.96) died, with a lethality rate of 55.08% in the studied population (CI 95% 49.20 - 51.15), which is considered high, and proves the severity of the disease. The mortality coefficient for sporotrichosis was 4.6 deaths for every 1000 cats (CI 95% 3.4 - 6 per 1000 cats) of the population of the Barreiro’s county. Animals with a positive diagnosis were 6.30 times more likely to die than negative animals (p< 0.05, OR 6.30, CI 95% 2,79-14,42).

Among the positive animals, 82.85% (CI 95% 82.10 - 83.58) showed an unfavorable progression of the disease and only 7.62% (CI 95% 7.12 - 8.16) were cured, after undergoing treatment. Notably, there were reports of deaths caused by both natural causes and by the actions of those responsible for the animals (such as poisoning or setting the animals on fire).

A number of factors may be related to the reduction in the cure of the disease, such as the high cost of medication, socioeconomically poor region, prolonged period of treatment, weak accountability of the cat owners, and an absence of a public policy that permits access to treatment and free veterinary monitoring of these animals.

In cases of death or euthanasia of the animals, we tried to find out what happened to the corpses. The percentage of positive cases that resulted in death and sent to incineration was 46.15% (CI 95% 33.70 - 58.97, 30/65). This result was possible because, at the time of the first visit, the owners were informed about the need to incinerate the corpse of the infected animals and free access to this service was provided by the UFMG. Despite this, a considerable percentage of dead positive animals (29.23%) (CI 95% 17.31 - 40.19, 19/65) were inappropriately discarded. Among these, 63.2% (CI 95% 50.20 - 74.72) were thrown in barren land, 31.2% (95% CI 28.40 - 34.14) were discarded into household trash, and one of the owners did not inform the exact location of disposal. Therefore, public policies of environmental education and actions that will emphasize the correct disposal of animal corpses with sporotrichosis are of great importance to prevent contaminating the environment, given the saprophytic nature of *Sporothrix* sp.

## Conclusion

According to previous literature, this is the first report on the epidemic of sporotrichosis in Minas Gerais. Sporotrichosis is present in the municipality of Belo Horizonte with wide distribution, affecting humans and cats, and a high mortality in domestic feline cases has been observed in the county of Barreiro. Suspected animals with free access to the streets have a greater chance of a positive result for sporotrichosis; therefore, public environmental education is urgently required to modify the relationship between cat owners and their animals. The disease presented an unfavorable prognosis for cats, even though a treatment exists. Additionally, inappropriate disposal of corpses was also seen. Therefore, the free offer for treatment and veterinary care to these animals should be taken into consideration, as well as the collection and incineration of the dead ones, as measures of public health, followed by the guidance and care for the human patient.

## Acknowledgments

We thank the DZ/SMSA and LAMICO for providing the data; veterinarians and public health professionals for the collection of samples and information with tutors; to veterinary residents of EV / UFMG; and EV by logistic support.

## Supporting information

**S1 Table - Distribution of household cats analyzed as per the demographic variables, county of Barreiro, Belo Horizonte, Minas Gerais, August 2016 to June 2018.**

**S2 Table - Distribution of household cats suspected of sporotrichosis analyzed, according to coverage areas, county of Barreiro, Belo Horizonte, Minas Gerais, August 2016 to June 2018.**

**S3 Table - Distribution of domestic felines analyzed as per the environmental variables.**

**S4 Table - Distribution of domestic cats collected according to the outcome and association of the variables with the positive diagnosis of sporotrichosis, in county of Barreiro, 2017/2018.**

